# Fibroblast-specific IL11 signaling is required for lung fibrosis and inflammation

**DOI:** 10.1101/801852

**Authors:** Benjamin Ng, Jinrui Dong, Sivakumar Viswanathan, Anissa A. Widjaja, Bhairav S. Paleja, Eleonora Adami, Nicole SJ. Ko, Mao Wang, Stella Lim, Jessie Tan, Sonia P. Chothani, Salvatore Albani, Sebastian Schafer, Stuart A. Cook

## Abstract

Tissue injury leads to activation of resident stromal, parenchymal and immune cells to initiate reparative processes that, if unresolved, can lead to fibrosis and organ damage. The directionality of effect between fibrosis and inflammation in the lung has been a point of debate for many years. Here, we tested the hypothesis that Interleukin 11 (IL11) signaling in fibroblasts is of primary importance for pulmonary fibrosis and that this event is upstream of lung inflammation. We generated mice with loxP-flanked *Il11ra1* alleles and crossed them to a *Col1a2*-CreERT strain to enable *Il11ra1* deletion in adult fibroblasts (*Il11ra1*-CKO mice). Lung fibroblasts from *Il11ra1*-CKO mice were selectively deleted for *Il11ra1* and refractory to TGFβ1 stimulation. In the mouse model of bleomycin-induced lung fibrosis, *Il11ra1*-CKO mice had markedly reduced pulmonary fibrosis and lesser lung damage, which was accompanied by diminished ERK activation in the stromal compartment. Bleomycin lung injury in *Il11ra1*-CKO mice was also associated with diminished STAT3 activation in inflammatory cells, fewer pulmonary immune cell infiltrates and almost complete inhibition of NF-kB activation. These data reveal an essential role for IL11 signaling in fibroblasts for lung fibrosis and show that inflammation in the lung can be secondary to stromal activation.

## INTRODUCTION

Fibrosis is a final common pathology underlying many chronic diseases and a major cause of morbidity and mortality (1). The main effector cells in fibrotic diseases are tissue-resident fibroblasts that are activated by local signals in injured tissues to become myofibroblasts. Myofibroblasts secrete large amounts of collagen, are pro-inflammatory and invade and disrupt tissue architecture, causing organ dysfunction (2, 3). There remains an unresolved issue as to the timing and directionality between tissue fibrogenesis and inflammation, notably in the lung and specifically in idiopathic pulmonary fibrosis (IPF).

Over decades, opinion has swung from the view that IPF is an inflammation-first-driven condition to a disease better characterized by prolonged epithelial injury, aberrant wound healing and, perhaps, a secondary wave of inflammation (4). In recent years, a primary immune role for Th2 polarized T cells and the Th2 cytokine IL13 gained favor in IPF fibrogenesis but targeting IL13 in clinical trials was ineffective (5). It remains to be resolved as to what are the key sentinel cells in the lung, immune and/or stromal, and how fibrosis and inflammation arise temporally: in series, in parallel or independently (6, 7).

IL11 is a member of the IL6 family of cytokines that share the use of the GP130 receptor as part of their multimeric receptor-ligand complexes for signal transduction. IL11 is secreted when tissue-resident fibroblasts are activated by profibrotic stimuli and binds IL11RA, which is highly expressed on stromal cells (e.g. fibroblasts and smooth muscle cells) (2). In recent studies, we documented an important role for IL11 in heart, kidney, lung and liver fibrosis (2, 3, 8) where it drives fibroblasts or hepatic stellate cells to become disease-causing myofibroblasts.

In mice, fibroblast-specific transgenic expression or systemic administration of IL11 induces fibroblast-to-myofibroblast transformation, causing widespread organ fibrosis, whereas mice with germline, global *Il11ra1* deletion are protected from fibrosis (2, 3, 8). Furthermore, inhibition of IL11 signaling using neutralizing antibodies prevents and reverses established fibrosis and inflammation in a mouse model of lung injury and in diet-induced non-alcoholic steatohepatitis (3, 8). Whilst these studies robustly annotate IL11 as a profibrotic factor that appears also important for inflammation, they do not address the directionality of these effects. We set out to dissect the fibroblast-specific effects of IL11 signaling to examine the relationship between fibroblast activation and inflammation in the lung.

Here we report the generation of a new *Il11ra1*-floxed mouse that we crossed to a *Col1a2*-CreERT strain to enable *Il11ra1* deletion in a temporal and conditional manner, specifically in adult fibroblasts. We subjected these mice to bleomycin (BLM) lung injury to explored the hypothesis that fibroblast-specific IL11 signaling is of primary importance for lung fibrosis and that IL11-mediated myofibroblast activation is required for and precedes lung inflammation.

## COMBINED RESULTS AND DISCUSSION

In our previous studies, we used the global *l11ra1* knockout mice to examine the role of IL11 signaling in various models of fibrotic disease. To further understand the cell-autonomous effects of IL11 signaling in fibrogenesis, we generated a *Il11ra1*-floxed mouse whereby *Il11ra1* could be conditionally deleted in a cell type-specific and temporal manner. Using CRISPR/Cas9 technique, we introduced loxP sites into the *Il11ra1* gene locus to allow for the conditional deletion of exons 4 to 7 (transcript Il11ra1-204: ENSMUST00000108042.2) (**Figure 1A** and **Supplementary Figure 1, A** and **B**). Upon Cre recombinase-mediated excision, it was predicted that the excision of exons 4 and 7 would lead to the generation of a premature stop codon and truncation of the Il11ra1 protein just after the N-terminal signal peptide sequence of *Il11ra1*, resulting in a null allele.

**Figure 1.**
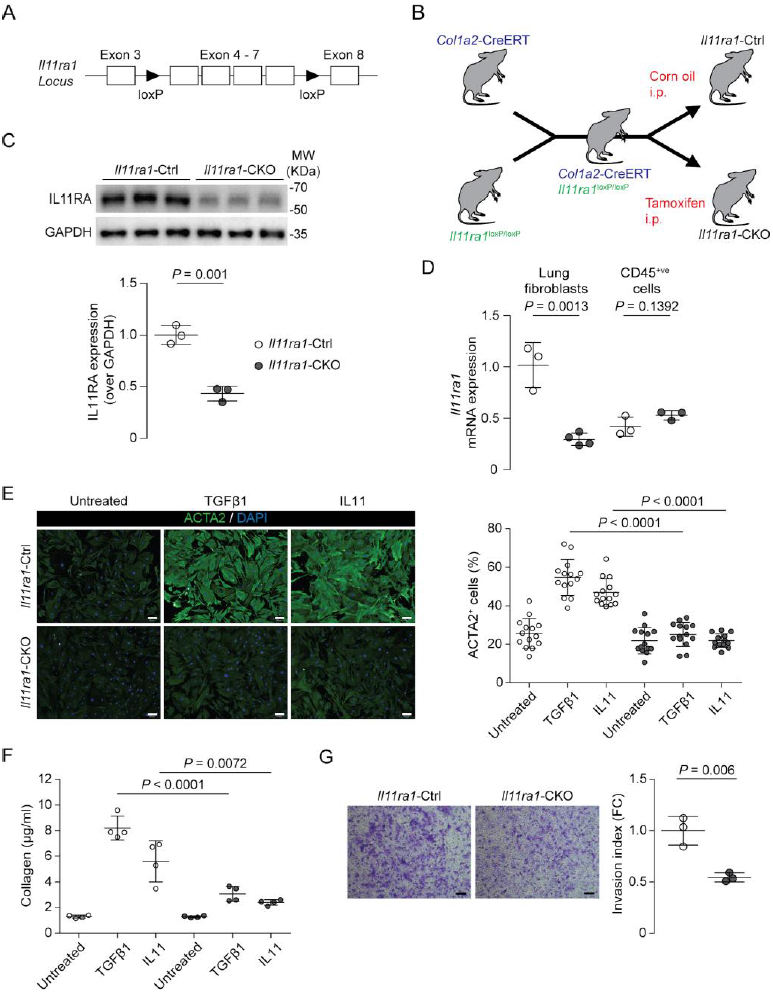
Generation of fibroblast-specific *Il11ra1*-deficient mice. (**A**) Schematic of loxP sites introduced into introns 3 and 7 of the mouse *Il11ra1* locus. (**B**) Schematic showing the generation of fibroblast specific-*Il11ra1* knockout (*Il11ra1*-CKO) mice and control (*Il11ra1*-Ctrl) mice. (**C**) Western blot of IL11RA expression in lung homogenates from *Il11ra1*-CKO and *Il11ra1*-Ctrl mice. (**D**) Gene expression of *Il11ra1* in fibroblasts and CD45-positive cells isolated from the lungs of *Il11ra1*-Ctrl and *Il11ra1*-CKO mice (*n* = 3-4). (**E**) Automated fluorescence imaging and immunofluorescence quantification of ACTA2^+^ cells from TGFβ1- or IL11-treated (5 ng/ml, 24h) lung fibroblasts from *Il11ra1*-CKO and *Il11ra1*-Ctrl mice. Cells were counterstained with DAPI to visualize the nuclei. One representative dataset from four independent experiments is shown (14 measurements / condition / experiment). Scale bars, 100 μm. (**F**) Total secreted collagen in the culture supernatant from **E** were quantified by Sirius Red collagen assay (*n* =4). (**G**) Matrigel invasion capacity of lung fibroblasts from *Il11ra1*-CKO and *Il11ra1*-Ctrl mice were determined (*n* = 3). Scale bars, 150 μm. Data shown are mean ± SD. *P* values were determined by Student’s *t*-test.

*Il11ra1*-floxed mutant mice were generated on the C57BL/6 background and mice homozygous for the floxed allele (*Il11ra1*^loxP/loxP^ mice) were born in normal Mendelian ratios and appear healthy from birth and during adult life. Female *Il11ra1*^loxP/loxP^ mice were fertile, as compared to germline, global *Il11ra1*-deleted female mice, which showed that the insertion of loxP sequences had no detrimental effects on *Il11ra1* gene function. Preliminary validation experiments were performed on fibroblasts isolated from the lungs of *Il11ra1*^loxP/loxP^ mice. After adenoviral delivery of Cre recombinase to *Il11ra1*^loxP/loxP^ fibroblasts, we confirmed a ∼80% knockdown of *Il11ra1* RNA as compared to cells transduced with control Adenovirus-GFP vectors (**Supplementary Figure 1C**).

Next, to determine if IL11 signaling in fibroblasts is important for lung fibrogenesis. We crossed homozygous *Il11ra1*^loxP/loxP^ mice to mice that express tamoxifen-inducible Cre recombinase driven by the mouse collagen type I alpha 2 promoter (*Col1a2*-CreERT mice), to generate *Col1a2*-CreERT *Il11ra1*^loxP/loxP^ mice (**Figure 1B** and **Supplementary Figure 2A**). To induce conditional knockdown of *Il11ra1* in fibroblasts, we injected 8 week old female *Col1a2*-CreERT *Il11ra1*^loxP/loxP^ mice with tamoxifen (referred to as *Il11ra1*-CKO mice), while littermates controls were injected with corn oil (referred to as *Il11ra1*-Ctrl mice) (**Figure 1B**). After five days of tamoxifen treatment, there was a 56% reduction in IL11RA protein levels in whole lung lysates from *Il11ra1*-CKO mice, which may be expected given the robust expression of *Il11ra1* on epithelial and vascular smooth muscle cells (**Figure 1C**) (9, 10). Furthermore, there was a 70% reduction of *Il11ra1* mRNA expression in primary fibroblasts isolated from the lungs of *Il11ra1*-CKO mice as compared to cells from *Il11ra1*-Ctrl mice (**Figure 1D**). In keeping with fibroblast-specific effects, tamoxifen treatment did not affect the expression of *Il11ra1* in CD45-positive cells from the lungs of *Il11ra1*-CKO mice.

**Figure 2.**
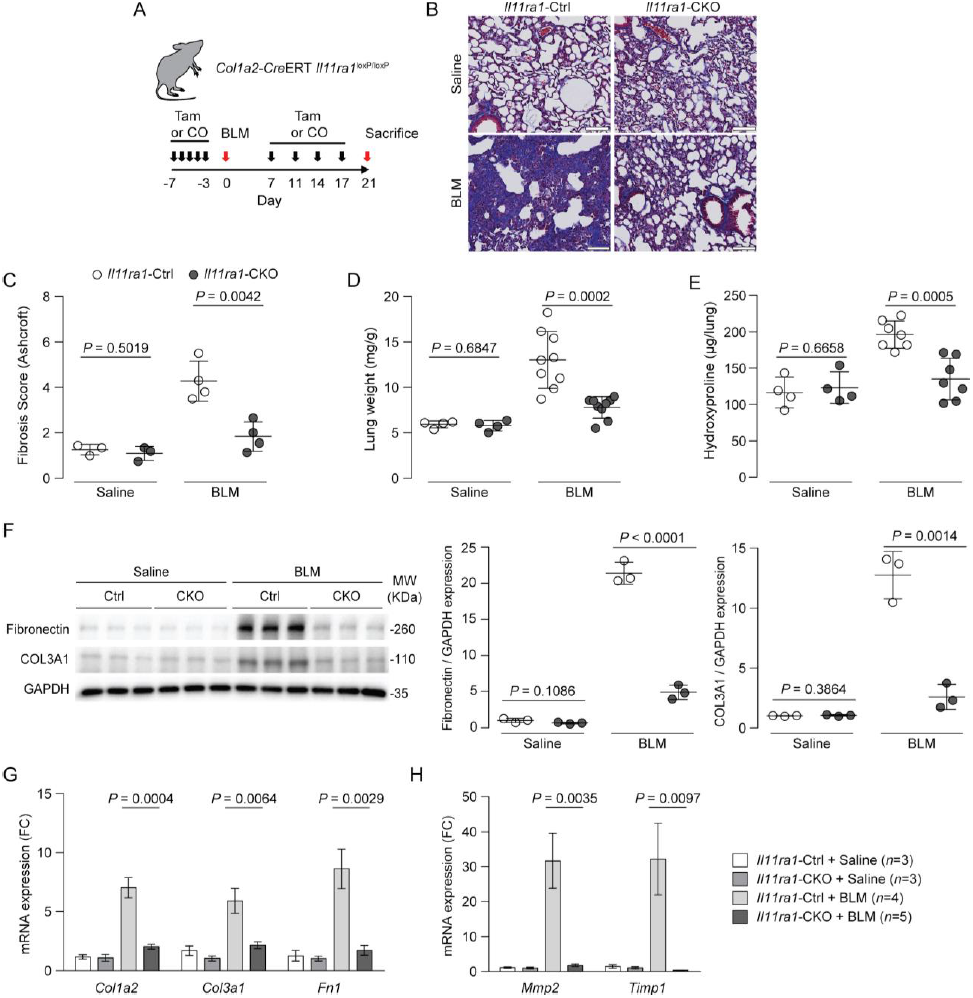
Fibroblast-specific deletion of *Il11ra1* prevents bleomycin-induced lung fibrosis. (**A**) Schematic showing the induction of pulmonary fibrosis in fibroblast specific-*Il11ra1* knockout mice. A single dose of bleomycin (BLM) was administered oropharyngeally and tamoxifen (Tam) or vehicle (corn oil: CO) was administered intraperitoneally at the indicated time points. (**B**) Masson’s trichrome staining of lung sections from *Il11ra1*-CKO or control mice 21d post-BLM. Scale bars, 100 μm. (**C**) Fibrosis score (*n* = 3-4), (**D**) lung weight to body weight indices and (**E**) lung hydroxyproline content of *Il11ra1*-CKO or control mice 21d post-BLM (*n* = 4-9). (**F**) Western blot and quantification of Fibronectin and COL3A1 expression in lung homogenates from *Il11ra1*-CKO or control mice 21d post-BLM (*n* = 3). (**G**-**H**) Expression of extracellular matrix and protease genes in lung lysates from *Il11ra1*-CKO or control mice 21d post-BLM (*n* = 3-5). Data shown in C-F are mean ± SD, and G and H are mean ± SEM. *P* values were determined by Student’s *t*-test.

Fibroblasts from mice with germline, global *Il11ra1* deletion (*Il11ra1*^*-/-*^) cannot upregulate fibrogenic proteins when stimulated with various fibrogenic stimuli, including TGFβ1 (2, 3). Thus, we examined the fibrogenic potential of TGFβ1 in lung fibroblasts isolated from *Il11ra1*-CKO mice. Lung fibroblasts were stimulated with recombinant mouse TGFβ1 (5 ng/ml; 24 hours) and activation monitored using automated high-throughput immunofluorescence imaging, collagen secretion and transwell matrigel invasion assays. Consistent with earlier studies, *Il11ra1*-CKO fibroblasts did not differentiate into myofibroblasts following TGFβ1 stimulation. Imaging analysis showed that ACTA2, COL1A1 expression and cell proliferation (as determined by EdU staining) were markedly decreased in TGFβ1-stimulated *Il11ra1*-CKO cells, as compared to controls (**Figure 1E** and **Supplementary Figure 2, B** and **C**). Collagen secretion into the culture medium was also greatly reduced from TGFβ1-treated *Il11ra1*-CKO fibroblasts (**Figure 1F**). Furthermore, recombinant mouse IL11 induced robust activation of *Il11ra1*-Ctrl fibroblasts, whereas the fibrotic response was greatly reduced in IL11-treated *Il11ra1*-CKO fibroblasts (**Figure 1, E** and **F**). Activated myofibroblasts acquire an invasive phenotype and we have shown that IL11 is crucial in the regulation of this process (3). We tested the ability of *Il11ra1*-CKO fibroblasts to invade ECM-coated Matrigel matrix, and found that *l11ra1*-CKO fibroblasts had reduced invasive capacity as compared to controls (**Figure 1G**).

Next, we determined the effect of fibroblast-specific *Il11ra1* deletion on the development of lung fibrosis using the bleomycin (BLM) model of pulmonary injury, which we characterised previously (3). *Col1a2*-CreERT *Il11ra1*^*fl/fl*^ mice mice were injected with five doses of tamoxifen prior to BLM challenge and administered an additional four additional doses post-BLM to maintain *Il11ra1* suppression in fibroblasts populations (**Figure 2A**). Histological examination of the lungs from saline-treated *Il11ra1*-CKO mice appeared normal. Following BLM injury (day 21), histology analysis revealed marked and highly significant reduction (84%, P=0.0042) in collagen accumulation and lung damage in *Il11ra1*-CKO mice as compared to controls (**Figure 2, B** and **C**). Consistent with the histology, lung weights and lung hydroxyproline content of BLM-treated *Il11ra1*-CKO mice were similarly significantly and profoundly decreased (reduction: 74%, P=0.0002; 82%, P=0.0005, respectively) as compared to controls (**Figure 2, D** and **E**). These changes were accompanied by notable reductions in Fibronectin and COL3A1 protein expression (**Figure 2F**) and a significant decrease in ECM and protease gene expression (*Col1a2, Col3a1, Fn1, Mmp2, Timp1*) (**Figure 2, G** and **H**) in lung lysates of *Il11ra1*-CKO mice.

The BLM model of lung injury is characterized by a strong initial inflammatory response which correlates with the extent of fibrosis at later time points (11). We evaluated whether the inactivation of *Il11ra1* in fibroblasts altered lung inflammatory responses and/or leukocyte recruitment following BLM. Western blot analysis of p-NF-kB, a pivotal mediator of inflammation across the spectrum of innate and acquired immune cells (12), revealed that *Il11ra1*-deletion in fibroblasts markedly reduced NF-kB activation in total lung lysates after BLM injury (**Figure 3A**). Gene expression analysis of *Il1b, Il6, Ccl2* and *Cxcl1* by qRT-PCR showed a reduction in the innate inflammatory response (**Figure 3B**). To examine further these anti-inflammatory effects, we studied leukocyte abundance in the lungs by immunostaining for CD45. This showed that *Il11ra1*-CKO mice had significantly reduced CD45^+ve^ immune cell lung infiltrates on day 21 post-BLM (**Figure 3C** and **Supplementary Figure 3A**).

**Figure 3.**
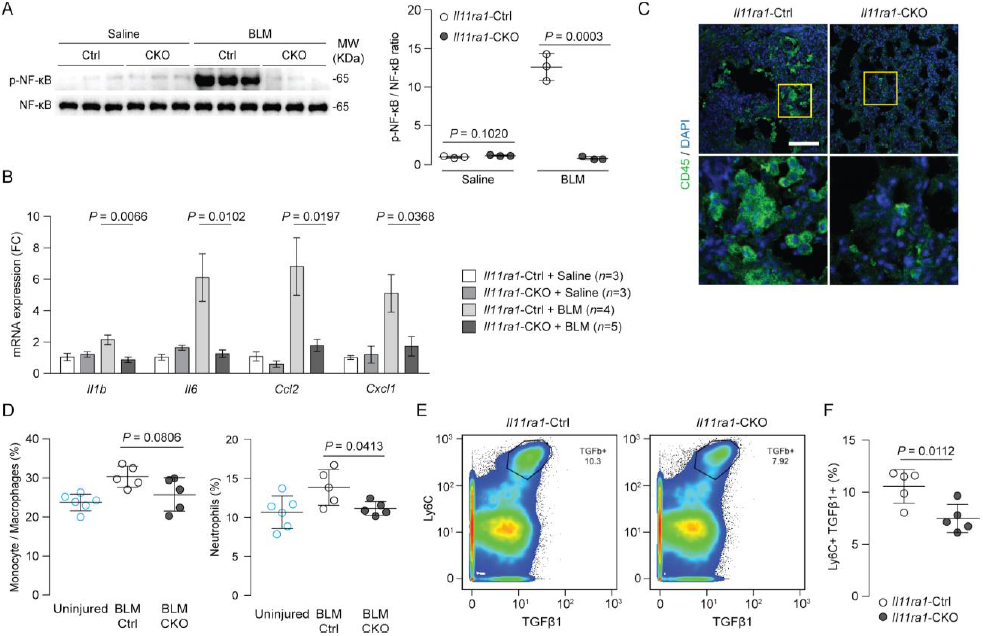
Fibroblast-specific deletion of *Il11ra1* inhibits bleomycin-induced lung inflammation. (**A**) Western blot of phosphorylated and total NF-κB expression in lung homogenates of *Il11ra1*-CKO or control mice 21d post-BLM (*n* = 3). (**B**) Expression of inflammatory response genes in lung lysates from *Il11ra1*-CKO or control mice 21d post-BLM (*n* = 3-5). Data shown are mean ± SEM. (**C**) Representative images of CD45 immunostaining in lung sections from *Il11ra1*-CKO or control mice 21d post-BLM. Scale bars, 100 μm. (**D**) CyTOF quantification of neutrophils, monocytes and macrophages in lung tissue from *Il11ra1*-CKO or control mice 14d post-BLM (n = 5-6). (**E**) Representative dot-plots and (**F**) quantification of Ly6C^+ve^TGFβ1^+ve^ cells in total CD45^+ve^ cells (following ex vivo stimulation with PMA) in lung tissue from *Il11ra1*-CKO or control mice 14d post-BLM (n = 5-6). Data shown in A, D and F are mean ± SD. *P* values were determined by Student’s *t*-test.

To examine the effects of *Il11ra1*-deletion in fibroblasts on what appeared to be downstream inflammatory responses in greater detail we used mass cytometry (CyTOF) (8, 13) to profile immune cell types in single cell suspensions of lung tissues from *Il11ra1*-Ctrl, -CKO mice and uninjured controls 14 days after BLM injury. As expected, we observed greater numbers of neutrophils, monocytes and macrophages in the lungs of BLM-treated *Il11ra1*-Ctrl mice as compared to uninjured controls. These cell populations were notably reduced in the lungs of BLM-treated *Il11ra1*-CKO as compared to *Il11ra1*-Ctrl (**Figure 3D**). We further profiled the infiltrating immune cell populations based on their pro-fibrotic potential and documented specific reductions in Ly6C^+ve^TGFβ1^+ve^ cells in PMA-stimulated CD45^+ve^ cells from the lungs of *Il11ra1*-CKO mice as compared to *Il11ra1*-Ctrl (**Figure 3, E** and **F**). These data, along with our previous findings (3), show that IL11 signaling in fibroblasts is both necessary and sufficient for lung inflammation in the BLM model of lung injury. This places stromal-driven inflammation at the apex of the inflammatory response in the fibrotic lung in the BLM model of lung injury.

We next determined the activation status (phosphorylation) of ERK and STAT3, both of which have been implicated in the pathogenesis of BLM-induced lung injury (3, 14). Western blot analysis of lung homogenates showed robust activation of ERK1/2 and STAT3 in BLM-challenged *Il11ra1*-Ctrl mice (**Figure 4A**). In contrast, *Il11ra1*-CKO mice showed significant reduction in both ERK1/2 and STAT3 activation after bleomycin (**Figure 4A**). Next, we used immunofluorescence and confocal imaging to localize the expression of activated ERK and STAT3 in BLM-injured lungs. In *Il11ra1*-Ctrl mice, we detected high levels of both phosphorylated ERK and STAT3 in fibrotic regions, which were reduced in lung sections from *Il11ra1*-CKO mice (**Figure 4B**). Interestingly, phosphorylated STAT3 was most highly expressed in cells of leukocyte morphology and less prominent in the stroma. Consistent with this, immunostaining for both CD45 and p-STAT3 revealed that p-STAT3 colocalized to leukocytes, and that the proportions of these double positive cells were reduced in the lungs of *Il11ra1*-CKO mice (**Supplementary Figure 3B**).

**Figure 4.**
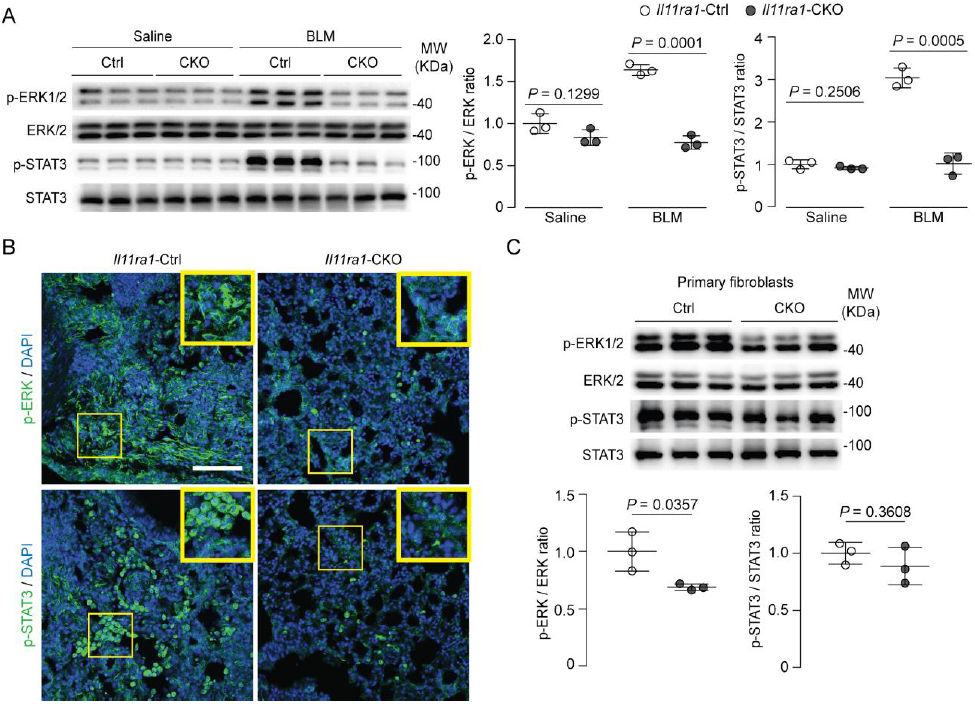
Reduced ERK and STAT3 activation in the lung and primary fibroblasts from bleomycin-treated mice with fibroblast-specific *Il11ra1* deletion. (**A**) Western blot and densitometry analysis of p-ERK1/2 and p-STAT3 expression in lung homogenates of *Il11ra1*-CKO or control mice 21d after BLM challenge (*n* = 3). (**B**) Images of p-ERK, p-STAT3 immunostaining in lung sections from *Il11ra1*-CKO or control mice 21d post-BLM. Scale bars, 100 μm. (**C**) Western blot and densitometry analysis of ERK1/2 and STAT3 expression in total cell lysates of primary fibroblasts isolated from *Il11ra1*-CKO or control mice 14d after BLM challenge (*n* = 3). Data shown are mean ± SD. *P* values were determined by Student’s *t*-test.

To determine if the reduction in the activation of ERK, STAT3 or NF-kB in lung lysates following fibroblast-specific *Il11ra1* deletion was due to lesser activation in fibroblasts, we isolated primary lung fibroblasts from *Il11ra1*-Ctrl and *Il11ra1*-CKO mice 14 days after BLM injury. Western blot analysis of primary fibroblast lysates showed a significant reduction in ERK activation in *Il11ra1*-CKO fibroblasts as compared to controls. In contrast, fibroblasts from BLM-treated *Il11ra1*-CKO mice had similar p-STAT3 and p-NF-kB levels when compared to *Il11ra1*-Ctrl (**Figure 3C** and **Supplementary Figure 3C**). Thus the overall reduction in p-STAT3 and p-NF-kB in the lungs of BLM-treated *Il11ra1*-CKO mice reflects reduced abundance of infiltrating inflammatory cells. Whereas, lower ERK activation in lung lysates from *Il11ra1*-CKO mice reflects lesser ERK activation in activated pulmonary fibroblasts.

We have previously identified a central role for IL11 signaling in fibrotic disease of the heart, liver and lung (2, 3, 8). Chronic disease is often characterized by localized inflammation and resident sentinel cells, which can be of mesenchymal or immune lineages, are important determinants of this tissue-specific immune response (6, 15). To better understand the contribution of IL11 signaling in specific cell types in disease pathology, we created the *Il11ra1*^loxP/loxP^ mouse for temporal and conditional inactivation of the *Il11ra1* gene.

Confirming our previous observations using global, germline *Il11ra1* deleted mice or neutralizing antibodies (3), adult pulmonary fibroblasts conditionally lacking IL11 signaling are unresponsive to pro-fibrotic stimuli *in vitro* (**Figure 1**). Histological scoring of lung damage and monitoring of RNA- and protein-based markers of pulmonary fibrosis revealed that *Il11ra1*-CKO mice are protected from BLM-induced lung fibrosis. These data define the critical role of IL11-dependent fibroblast activation in pathogenic ECM deposition in the BLM lung fibrosis model (**Figure 2**).

While fibroblasts have been implicated in inflammation-associated pathology of rheumatoid arthritis, colitis and IPF (7, 15–18), it has not been shown conclusively whether the stroma is a modifier or primary driver of the inflammatory response. In the BLM model of lung fibrosis, we found that the deletion of IL11 signaling in fibroblasts alone is sufficient to protect mice from inflammation in the lung. Remarkably, following BLM injury *Il11ra1*-CKO mice showed minimal activation of the NF-kB pathway, which is the prototypical activator of both the innate and acquired immune system (12), whereas control mice had marked induction of p-NF-kB. Other pro-inflammatory markers, such as *Il1b* or *Il6* were also suppressed in *Il11ra1*-CKO mice following BLM. *Il11ra1*-deletion in fibroblasts also reduced the expression of chemokines such as *Ccl2* and *Cxcl1* and blocked pulmonary infiltration of Ly6C^+ve^ neutrophils, monocytes and macrophages (**Figure 3**). Ly6C^+ve^ macrophages have been shown to drive fibrosis, in-part via their interactions with stromal cells (19–21). Similarly, neutrophilic infiltration contributes to BLM-induced lung fibrosis via their release of neutrophil elastase, which promotes TGFβ activation and myofibroblast differentiation (22–25). Our observations suggest that stromal cells are not only activated by immune cells, but also of central importance for their recruitment. Deactivation of the stroma via IL11 neutralization can break the vicious cycle of pathologic stroma- and immune-cell crosstalk and prevent both fibrosis and inflammation in IPF.

Western blotting of whole lungs revealed that both STAT3 (viewed as canonical downstream of GP130), as well as non-canonical ERK signaling, were reduced in the BLM damaged lungs of *l11ra1*-CKO mice (**Figure 4A**). This is perhaps surprising, as we genetically block IL11 signaling in fibroblasts only in *Il11ra1*-CKO mice and we have also shown previously that IL11 at physiological levels primarily activates ERK, but not STAT3, signaling in fibroblasts *in vitro* (2, 3, 8).

To identify the cellular compartment(s) characterized by ERK and/or STAT activation, we performed immunostaining of lung sections using p-ERK and p-STAT3 antibodies. In BLM-treated control animals, ERK activation was detected in clusters of spindle-shaped fibroblast-like cells. In contrast, STAT3 activation was localized to a different, round-shaped, cell-type, which we showed to be CD45^+ve^ immune cells (**Supplementary Figure 3**). In the *l11ra1*-CKO mouse, ERK activation was reduced in the stromal compartment, which we confirmed *ex vivo* in primary fibroblast cultures (**Figure 4C**). We suggest this is likely a direct effect of IL11 inhibition on this cell type. Immunostaining of lung sections showed STAT3 activity in immune cells still occurred in the *Il11ra1*-CKO but that number of CD45^+ve^ cells was greatly reduced. Thus the anti-inflammatory properties of IL11 neutralization appear to be indirect and reflect a less activated stroma (**Figure 3**). Western blotting of the whole lung lysates thus shows a reduction in p-STAT3^+ve^ cells and does not indicate a reduction of STAT3 activation in stromal or parenchymal cells.

Our data support the growing concept that fibroblasts are immunologically active and show that *Il11ra1*-dependent fibroblast activation can lead to inflammation (6). Further studies are required to delineate whether distinct subpopulations of IL11-responsive fibroblasts drive lung fibrosis and inflammation. It will also be important to examine whether IL11-dependent, fibroblast-mediated inflammation occurs in other lung diseases, such as asthma (26, 27) or additional conditions such as rheumatoid arthritis or colitis (15–18). Preliminary functional studies suggest this may be the case, at least for colitis (10), which is characterized by IL11-expressing *inflammatory fibroblasts* that predict disease severity in patients (17).

## METHODS

### Study approval

Animal studies were performed in accordance to approved protocols by the SingHealth Institutional Animal Care and Use Committee (IACUC).

Detailed methods are provided in the Supplementary Material.

## AUTHOR CONTRIBUTIONS

B.N. and S.A.C designed the study. B.N., J.D., N.S.K., and J.T. performed *in vivo* studies. B.N., J.D., S.V., A.A.W, M.W. and S.L. performed *ex vivo* and *in vitro* studies. B.S.P. and S.A conducted mass cytometry experiments. B.N., S.V., B.S.P., E.A. and S.P.C. analyzed the data. B.N., S.S. and S.A.C wrote the manuscript.

## ACKNOWLEDGMENTS

The authors would like to acknowledge B.L.George for his technical expertise and support. The research was supported by the National Medical Research Council (NMRC) Singapore STaR awards to S.A.C. (NMRC/STaR/0011/2012 and NMRC/STaR/0029/2017), the NMRC Centre Grant to the NHCS, Goh Foundation, Tanoto Foundation and a grant from the Fondation Leducq.

## SUPPLEMENTAL METHODS

### Animal models

#### Generation of Il11ra1 floxed mice

CRISPR/Cas9 technique was used to introduce loxP sequences into the *Il11ra1* gene (ENSMUSG00000073889; Transcript Il11ra1-204: ENSMUST00000108042.2) to allow for the conditional deletion of exons 4 to 7, which will result in a null allele upon Cre recombinase-mediated excision. Single guide RNAs (sgRNAs) with recognition sites on introns 4 and 7 along with Cas9 and a targeting construct containing two loxP sequences were microinjected into fertilized zygotes, and subsequently transferred into pseudopregnant mice (Shanghai Model Organisms Center, Inc). Insertion of loxP sites into the *Il11ra1* gene locus were verified by sequencing. Mutant Il11ra1-floxed offspring were generated on a C57BL/6 background and identified by genotyping to detect the insertion of a loxP site in intron 7 using the following primers: 5’-CATTACCTACACATCCTTCCC-3’ and 5’-CCTCCCACACTCAGATTAAGA-3’.

#### Generation of fibroblast-specific Il11ra1 knockout mice

To direct fibroblast-specific *Il11ra1* deletion in mice, homozygous *Il11ra1*-floxed mice (*Il11ra1*^loxP/loxP^) were crossed with hemizygous *Col1a2*-CreERT mice (28) (Jackson’s Lab) to generate Col1a2-CreERT *Il11ra1*^loxP/loxP^ progenies. Six to eight week old *Col1a2-Cre*ERT; *Il11ra1*^loxP/loxP^ mice received intraperitoneal injections of tamoxifen (50 mg kg^-1^ body weight) per day for 5 consecutive days before bleomycin challenge, followed by subsequent tamoxifen injections at day 7, 11, 14 and 17 after bleomycin challenge to delete *Il11ra1* from fibroblast populations (*Il11ra1*-CKO mice). Littermate animals were injected with corn oil as vehicle as controls (*Il11ra1*-Ctrl mice).

#### Bleomycin model of lung injury

Mice were anesthetized by isoflurane inhalation and bleomycin (Sigma-Aldrich) was administered oropharyngeally at a dose of 1 mg kg^-1^ body weight or saline as sham controls.

### Antibodies and reagents

Antibodies: Anti-ACTA2 (ab7817, abcam), anti-COL1A1 (ab34710, abcam), anti-COL3A1 antibody (sc-271249, Santa Cruz), anti-Fibronectin antibody (ab2413, Abcam), anti-p-ERK1/2 (4370, Cell Signaling), anti-ERK1/2 (4695, Cell Signaling), anti-p-STAT3 (4113, Cell Signaling), anti-STAT3 (4904, Cell Signaling), anti-GAPDH (2118, Cell Signaling), anti-CD45 (ab10558, abcam), p-NF-kB (3033, Cell Signaling), NF-kB (8242, Cell Signaling) and anti-IL11RA (130920, Santa Cruz). Recombinant proteins: TGFβ1 (7666-MB, R&D Systems), PDGF (220-BB, R&D Systems) and recombinant mouse IL11 (GenScript).

### Primary lung fibroblast cultures

Primary mouse lung fibroblasts were isolated from 8-12 week old *Il11ra1*-CKO and *Il11ra1*-Ctrl mice after 5 consecutive doses of tamoxifen (1 mg/kg) or corn oil respectively. Lungs were minced, digested for 30 minutes in DMEM containing 100 U ml^-1^ penicillin, 100 μg ml^-1^ streptomycin and 0.14 Wunsch U ml^-1^ Liberase (Roche) with mild agitation. Lung tissue were washed and subsequently cultured in complete DMEM supplemented with 15% FBS, 100 U ml^-1^ penicillin and 100 μg ml^-1^ streptomycin at 37°C. Fibroblasts were enriched via negative selection with magnetic beads against mouse CD45 (leukocytes), CD31 (endothelial) and CD326 (epithelial) using a QuadroMACS separator (Miltenyi Biotec) according to the manufacturer’s protocol. Mouse lung fibroblasts were used for downstream experiments between passages 3 to 5. To induce Cre recombinase expression, *Il11ra1*^loxP/loxP^ fibroblasts were transduced with Adenovirus-Cre-GFP (VectorBiolabs) or Adenovirus-CMV-GFP as control vector. One day after transduction, cells were lysed and mRNA expression of *Il11ra1* were analysed by RT-qPCR.

### In vitro immunofluorescence and image analysis

Immunofluorescence imaging and quantification of fibroblast activation were performed on the Operetta High Content Imaging platform (PerkinElmer) as previously described (2). Briefly, lung fibroblasts were seeded in 96-well CellCarrier plates (6005550, PerkinElmer) at a density of 5×10^3^ cells per well and following experimental conditions, cells were fixed in 4% paraformaldehyde (28908, Life Technologies) and permeabilized with 0.1% Triton X-100 in PBS. EdU-Alexa Fluor 488 was incorporated using a Click-iT Edu Labelling kit (C10350, Life Technologies) according to manufacturer’s protocol. Cells were blocked using 0.5% BSA and 0.1% Tween-20 in PBS before incubation with primary antibody (anti-ACTA2 or anti-COL1A1) and visualized using Alexa Fluor 488-conjugated secondary antibody (ab150113, Abcam). Cells were counterstained with DAPI (D1306, Life Technologies) in blocking solution. Plates were scanned and images were collected with an Operetta high-content imaging system 1483 (PerkinElmer). Each experiment was run in duplicate wells, 7 fixed fields per well were measured and data points from duplicate wells were combined for a total of n = 14 measurements per condition. The percentage of activated fibroblasts (ACTA2^+^ cells) and EdU^+^ cells in each field was quantified using the Harmony software version 3.5.2 (PerkinElmer). The measurement of COL1A1 fluorescence intensity was done using Columbus software version 2.7.1 (PerkinElmer). Fluorescence intensity for COL1A1 staining was normalized to the cell area detected in each field. Representative results using primary cells of one individual mouse are shown in the manuscript as described in the figure legends.

### Matrigel invasion assay

The invasive capacity of lung fibroblasts isolated from *Il11ra1*-CKO and *Il11ra1*-Ctrl mice were assayed using Boyden chamber invasion assay (Cell Biolabs Inc.). Equal numbers of fibroblasts in serum-free DMEM were seeded in duplicates onto ECM-coated matrigel and allowed to invade towards DMEM containing 2% FBS and PDGF (20 ng ml^-1^) as chemoattractants. After 24h of incubation at 37°C, media was removed and non-invasive cells were removed using cotton swabs. The cells that migrated or invaded towards the bottom chamber were stained with cell staining solution (Cell Biolabs Inc.). Invasive cells from 5 non-overlapping fields of each membrane were imaged and counted under 40x magnification.

### Histology

Freshly dissected mouse lungs were fixed in 10% Neutral buffered formalin for 24h, embedded in paraffin and sectioned for staining. For Masson’s trichrome staining of lung sections were described previously (3). Fibrosis scores in Masson’s trichrome stained lung sections were assessed using the Ashcroft score (29), and histopathological scoring was performed blinded to genotype and bleomycin treatment. For immunostaining of lung tissues, mouse lungs were firstly fixed overnight in 4% paraformaldehyde (Thermofisher) at 4°C, dehydrated in 20% sucrose in PBS for 72 hours and frozen in Optimal cutting temperature compound (OCT) prior to sectioning. Fifteen micrometer cryosections were first permeabilized with 0.1% Triton X-100 for 30 minutes and then blocked in 3% bovine serum albumin in PBS for 30 minutes. Primary antibodies concentrations were then applied according to manufacturer’s instructions and incubated at 4°C overnight, followed by secondary antibody incubation for 30 minutes at room temperature (anti-mouse or rabbit Alexa Fluor-555 and -647, Molecular Probes). Sections were mounted with ProLong glass antifade mountant (Molecular probes) and fluorescence images were acquired using a confocal laser scanning microscope (LSM 710, Zeiss).

### Colorimetric assays

For the detection of secreted collagen by cells in culture, the supernatant was first concentrated using polyethylene glycol solution (90626 Chondrex) prior to quantification using a Sirius red collagen detection kit (9062, Chondrex), according to manufacturer’s protocol. Total hydroxyproline content in the right lungs of mice were measured using a Quickzyme Total Collagen assay kit (Quickzyme Biosciences) according to the manufacturer’s protocol.

### RT-qPCR

Total RNA was extracted from snap-frozen mouse lung tissues using Trizol reagent (Invitrogen) followed by RNeasy column (Qiagen) purification and cDNA was prepared using an iScript cDNA synthesis kit (Biorad) following manufacturer’s instructions. Quantitative RT–PCR gene expression analysis was performed with TaqMan (Applied Biosystems) technology or with QuantiFast SYBR Green PCR kit (Qiagen) using a StepOnePlus (Applied Biosystem). Expression data were normalized to *Gapdh* mRNA expression using the 2^**-ΔΔCt**^ method to calculate the fold change. TaqMan probes were obtained from Thermo Fisher Scientific (*Col1a1*, Mm00801666_g1; *Col1a2*, Mm00483888_m1; *Col3a1*, Mm01254476_m1; *Mmp2*, Mm00439498_m1; *Timp1*, Mm01341361_m1; *Gapdh*, Mm99999915_g1). Inflammatory response genes primer sequences are as follows: *Cxcl1*, 5’-ACCCGCTCGCTTCTCTGT-3’ and 5’-AAGGGAGCTTCAGGGTCAAG-3’; *Ccl2* 5’-GAAGGAATGGGTCCAGACAT-3’ and 5’-ACGGGTCAACTTCACATTCA-3’; *Il6* 5’-CTCTGGGAAATCGTGGAAAT-3’ and 5’-CCAGTTTGGTAGCATCCATC-3’; *Il1b* 5’-CACAGCAGCACATCAACAAG-3’ and 5’-GTGCTCATGTCCTCATCCTG-3’. Deletion of *Il11ra1* was confirmed in lung fibroblasts isolated from tamoxifen-treated *Col1a2-Cre*ERT; *Il11ra1*^loxP/loxP^ mice using the following primers specific to Exon 4 of *Il11ra1*: 5’-CCCACCCGCTACCTTACTTC-3’ and 5’-CATGGACCACACATCGGGAG-3’.

### Immunoblotting

Western blot analysis was carried out on total protein extracts from snap-frozen mouse lung tissues. Tissues were homogenized in lysis buffer (RIPA buffer containing protease and phosphatase inhibitors (Roche) followed by centrifugation to clear the lysate. Equal amounts of protein lysates were separated by SDS-PAGE, transferred to a PVDF membrane, and incubated overnight with primary antibodies. Proteins were visualized using the ECL detection system (Pierce) with the appropriate secondary antibodies: anti-rabbit HRP (7074, Cell Signaling) or anti-mouse HRP (7076, Cell Signaling).

### Mass cytometry by Time of Flight (CyTOF)

Lung tissues were minced and digested with 100 µg/ml Collagenase IV and 20U/ml DNase I, at 37°C for 60 mins. Following digestion, cells were passed through 100 µm strainer to obtain single cell suspension. Cells were cryopreserved until further use. CyTOF staining was performed as previously described (13). For immunophenotyping by CyTOF, a panel of 36 antibodies that encompassed a broad range of immune cells types was designed. Briefly, cells were thawed, stained with cisplatin (Fluidigm) to identify live cells, followed by staining with metal-conjugated cell surface antibodies. Cells were then barcoded using 20-plex Palladium barcoding kit (Fuidigm) according to manufacturer’s protocol. After barcoding cells were subjected to intracellular antibody staining (TGFβ1). Cells were labeled with DNA intercalator before acquisition on Helios mass cytometer (Fluidigm). For analysis, live single cells were identified, followed by debarcoding to identify individual samples. Manual gating was performed using Flowjo software (Flowjo, LLC, USA) to identify different immune cell subsets.

### Statistics

Statistical analysis was performed using GraphPad Prism software (version 6.07), using two-sided Student’s t-tests. *P* values < 0.05 are regarded as statistically significant.

**Supplementary Figure 1.**
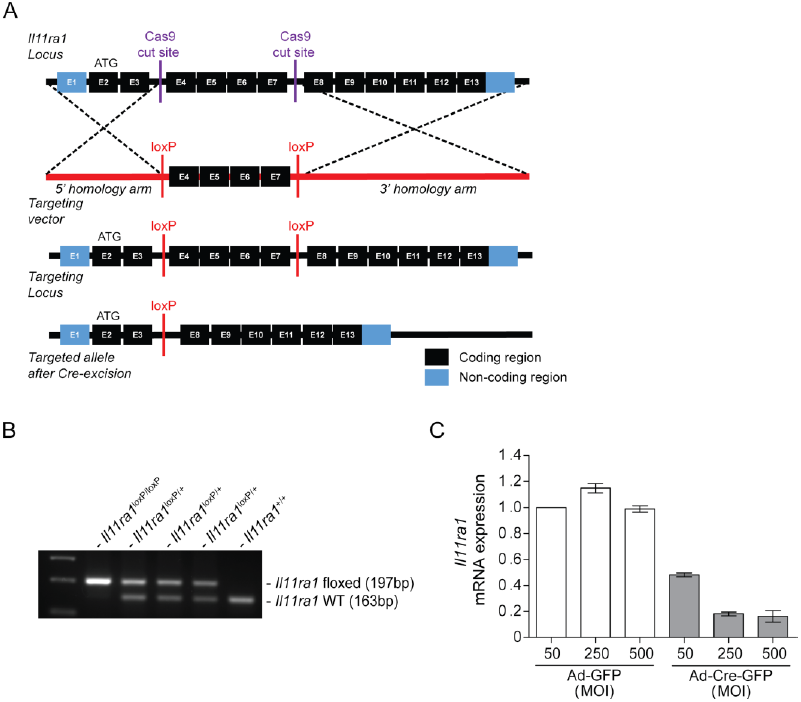
Schematic and genotyping of *Il11ra1* conditional knockout alleles. (**A**) Schematic design of CRISPR/Cas9 mediated insertion of loxP sites into introns 3 and 7 of the mouse *Il11ra1* locus (Transcript Il11ra1-204: ENSMUST00000108042.2). (**B**) PCR genotyping of mice with *Il11ra1*-floxed and wildtype alleles. (**C**) Conditional knockdown of *Il11ra1* mRNA was assessed by qRT-PCR in Adenovirus-Cre transduced lung fibroblasts isolated from *Il11ra1*^*loxP/loxP*^ mice (*n* = 2). MOI: multiplicity of infection.

**Supplementary Figure 2.**
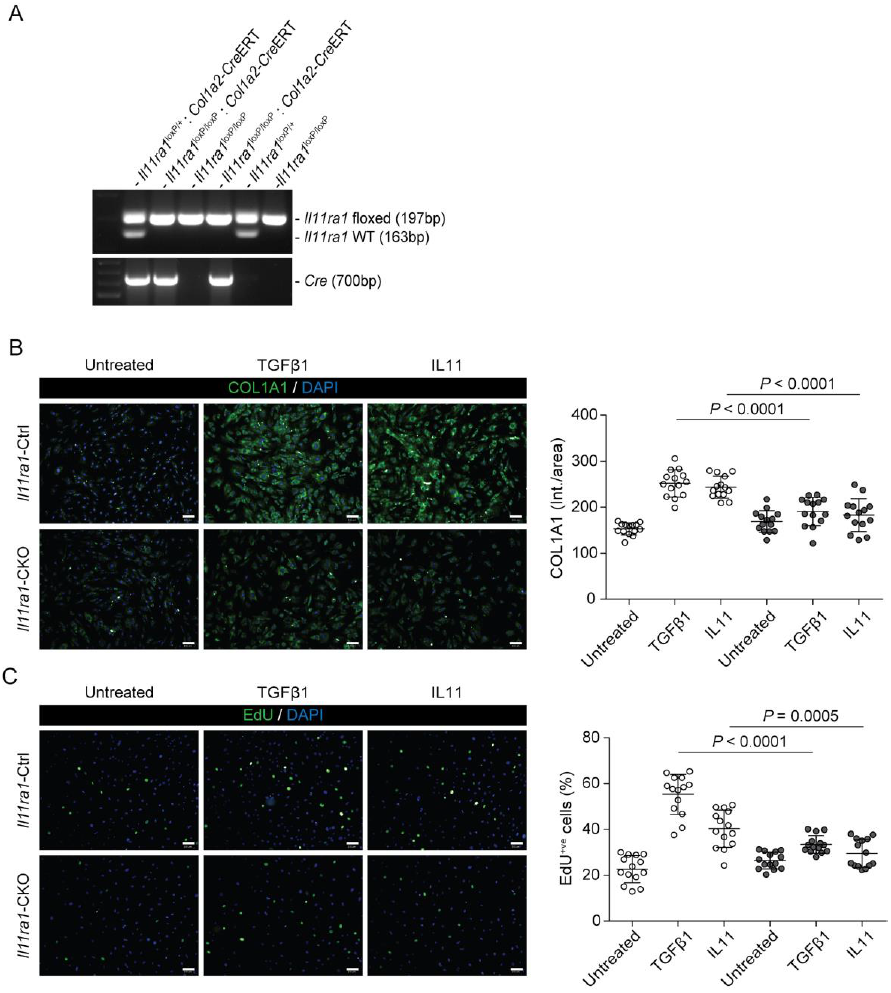
Collagen expression and cell proliferation are inhibited in TGFβ1-stimulated lung fibroblasts from fibroblast-specific *Il11ra1*-deficient mice. (**A**) PCR genotyping showing *Il11ra1* WT (163bp) *Il11ra1*-floxed (197bp) and *Cre* (700bp) PRC products from a cross between *Il11ra1*^loxP/+^ and *Col1a2*-CreERT *Il11ra1*^loxP/loxP^ mice. (**B**-**C**) Automated fluorescence imaging and immunofluorescence quantification of (**B**) COL1A1 expression (intensity/area) and (**C**) EdU^+ve^ cells in TGFβ1- or IL11-treated lung fibroblasts from *Il11ra1*-CKO or control mice (5 ng/ml, 24h). One representative dataset from four independent biological experiments is shown (14 measurements per condition per experiment). Scale bars, 100 μm. Data shown are mean ± SD. *P* values were determined by Student’s *t*-test.

**Supplementary Figure 3.**
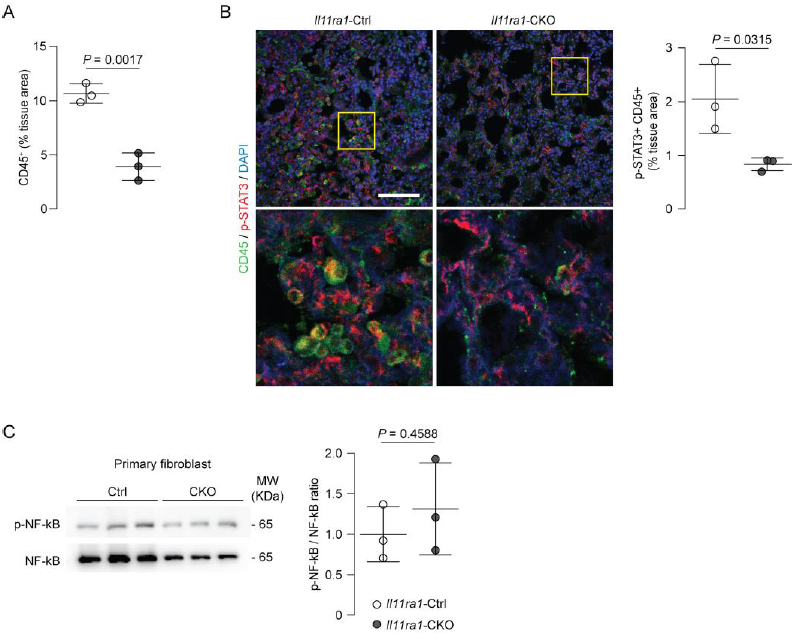
Normal expression of p-NF-kB in lung fibroblasts but reduced STAT3 activation in immune cells from the lungs of BLM-treated *Il11ra1*-CKO mice. (**A**) Quantification of CD45 positive cells in immunofluorescence images of lung sections from *Il11ra1*-CKO or control mice 21d post-BLM, depicted in **Figure 3**. (**B**) Representative images and quantification of CD45 and p-STAT3 immunostaining in lung sections from *Il11ra1*-CKO or control mice 21d post-BLM. Scale bars, 100 μm. (**C**) Western blot and densitometry analysis of p-NF-kB and NF-kB expression in total cell lysates of primary fibroblasts isolated from *Il11ra1*-CKO and control mice 14d after BLM challenge (*n* = 3). Data shown are mean ± SD. *P* values were determined by Student’s *t*-test.

## Notes

**Conflict of interest declaration:** S.A.C. and S.S. are co-inventors of the patent applications (WO2017103108, WO2017103108 A2, WO 2018/109174 A2, WO 2018/109170 A2) for “Treatment of fibrosis”. A.A.W., S.S., and S.A.C are co-inventors of the patent applications GB1900811.9, GB 1902419.9, GB1906597.8 for “Treatment of hepatotoxicity, nephrotoxicity, and metabolic diseases”. S.A.C. and S.S. are co-founders and shareholders of Enleofen Bio PTE LTD, a company (which S.A.C. is a director of) that develops anti-IL11 therapeutics.

